# Epidemiological Analysis of Ear Diseases in 221 Dogs in Northwest of China

**DOI:** 10.1101/541516

**Authors:** F-L. Tang, H-Q. Yang, X-W. Ma, D-Z. Lu

**Affiliations:** Department of Clinical Veterinary Medicine, College of Veterinary Medicine, Northwest A&F University, Yangling, Shaanxi 712100, P.R. China; Xi’an Teaching Hospital, Northwest A&F University, Xi’an, Shaanxi 71007, P.R. China; Department of Clinical Veterinary Medicine, College of Veterinary Medicine, China Agricultural University, Beijing 100193, P.R. China

**Keywords:** dog, ear diseases, otitis externa, epidemiology

## Abstract

Ear disease is a relatively common disease in veterinary, and it will great effect on the pets’ life. This study mainly collected and collated the cases of Xi’an Teaching Hospital of Northwest A&F University from 2012 to 2016, and conducted an epidemiological analysis of the incidence of ear disease in dogs. In this study, the common ear diseases in dogs were divided into otitis externa, otitis media, otitis interna, and ear hematoma. The results showed that the total number of dog ear disease cases was 221, the highest incidence was otitis externa (84.62%), and the main cause of otitis externa was the bacterial infection (44.10%), which followed by fungal infection (32.31%). The ratio of male to female is 1:1.5. In terms of the dog species, the highest incidence was Teddy (18.55%), followed by the Golden Retriever (10.41%). The most common ear diseases in puppies was otitis externa, ear canal hyperplasia and ear canal tumors occur mainly in older dogs. Through the analysis of the 221 cases of ear diseases, we could better understand the development and epidemiology of dog ear diseases, and provide some reference for clinical diagnosis, treatment and prevention of dog ear diseases.

In recent years, with the continuous improvement of people’s quality of life, more and more people choose to keep pets to accompany themselves, and the number of pets in Xi’an has also increased sharply. However, ear disease is one of common diseases in dogs, their characteristics include pruritus, long treatment period and easy to relapse. If the treatment is not timely, there may be secondary otitis media and otitis interna, which may affect the hearing of dogs. In severe cases, neurological symptoms may occur, which may have a bad impact on the life and health of dogs. Now, we have collected and sorted out a total of 221 cases of ear diseases in dogs from Xi’an Teaching Hospital of Northwest A&F University from 2012 to 2016, and classified them into four directions: otitis externa, otitis media, otitis interna and ear hematoma for epidemiological analysis, so as to provide references for the clinical diagnosis and treatment of ear diseases in dogs in the future.

## 1 Materials and Methods

### 1.1 Source of cases

The situation of dog ear diseases in Xi’an Teaching Hospital of Northwest A&F University from 2012 to 2016 were statistically analyzed by collecting cases, looking through historical cases, returning to telephone to obtain medical history data and laboratory results.

### 1.2 Inspection of body surface

Examining systematically the skin of the animals’ body surface, and observing carefully the ears of dogs for depilation, callus, dandruff, erythema and thickening, whether the ear canal is moist, with or without secretions and odor.

### 1.3 Collection of disease material

#### 1.3.1 Auricular skin scraping test

Firstly, cutting the coat at the junction of the affected area and the healthy area, and then the skin was gently scraped along the direction of the hair with a blunt blade until there was a slight bleeding. The scraped disease materials were placed on the slide, some of the disease materials were added with Rayleigh’s dye to observe if the bacteria exist in the microscopic examination. Another part of the disease materials were added with 2 drops of 10% KOH, and the cover glasses were slightly pressed. Heating it slightly, after the samples were transparent, observing if the insects existed by microscopy.

#### 1.3.2 Examination of ear canal secretion

Directly using sterile cotton swab to collect ear canal secretion, smearing the swab with samples on the slides. After dried, fixed and stained, observing them by microscopy.

### 1.4 Instrument inspection

Using the ear endoscope to check the condition of the ear canal, inserting the probe into the ear canal to see if there is any foreign body and the integrity of the middle ear structure, or doing biopsy of the ear canal tumor.

## 2 Results and Analysis

Counting the number of cases of dog ear diseases from 2012 to 2016, knowing the proportion of total cases of dogs with ear diseases, the prevalence and seasonality of ear diseases, the relationship between ear diseases and environment, age, variety, gender. It was analyzed to summarize the epidemiology of ear diseases.

### 2.1 Investigation results of the incidence of canine ear diseases

This study was conducted a statistical survey about outpatient medical records of Xi’an Teaching Hospital of Northwest A&F University from January 2012 to December 2016. The total number of outpatient cases in dogs was 20404, there was 221 dog ear diseases, accounting for 1.08%. (Table 1).

**Table 1.** The incidence of dog ear diseases from 2012 to 2016

| Year | 2012 | 2013 | 2014 | 2015 | 2016 |
| --- | --- | --- | --- | --- | --- |
| Total cases | 3224 | 4417 | 4196 | 4563 | 4004 |
| Number of dog<br>ear diseases | 38 | 34 | 48 | 51 | 50 |

### 2.2 Investigation results of the cause of dog ear diseases

Among the 221 ear diseases, the incidence of otitis externa was the highest, accounting for 84.62%; followed by ear hematoma, accounting for 6.33%; the incidence of otitis media and otitis interna respectively was 3.62% and 1.81%. The distribution of different types of dog ear diseases was shown in Figure 1.

**Fig. 1.**
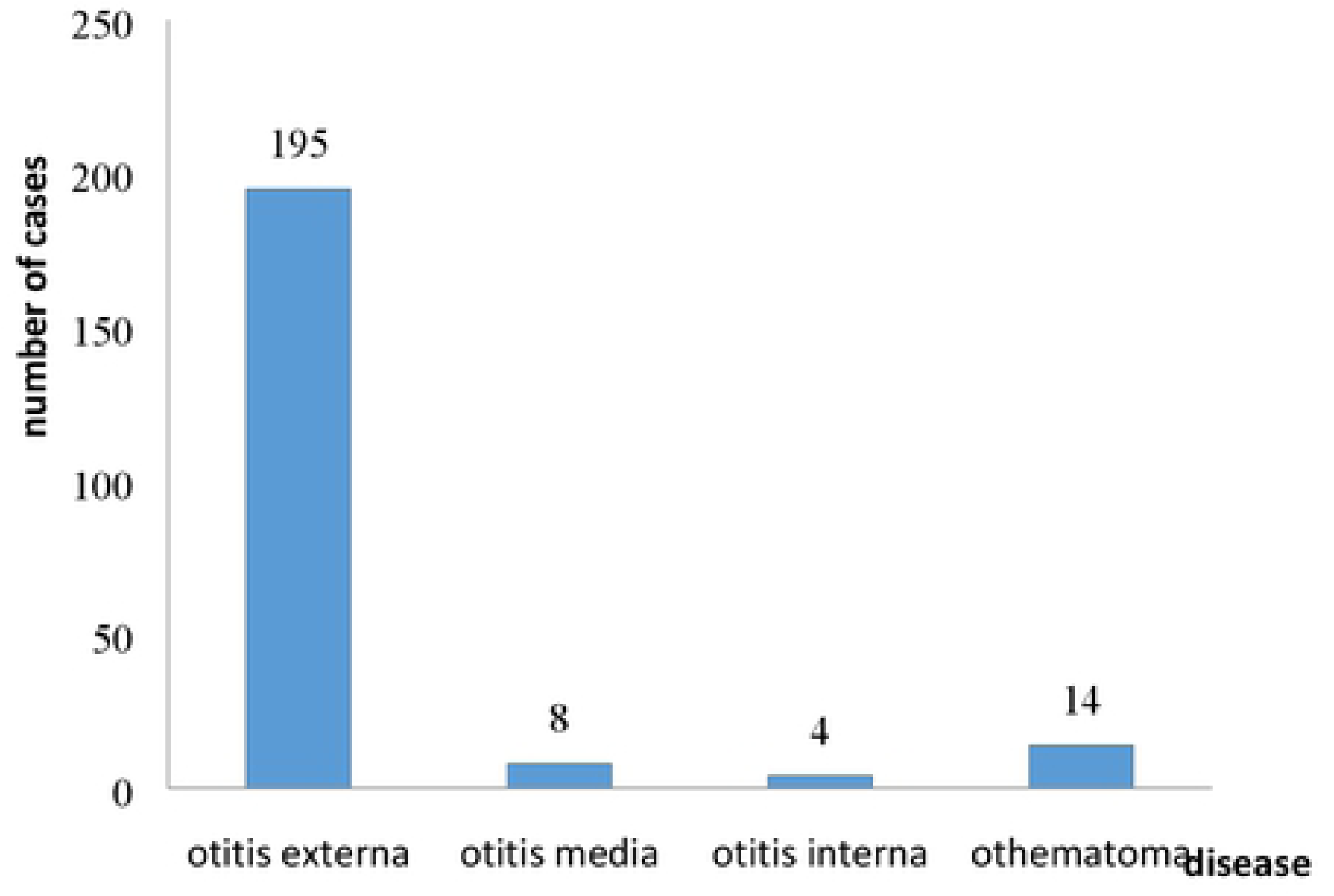
The distribution of different types of eardiseases from 2012 to 2016

In 195 cases of canine otitis externa, the incidence of bacterial otitis externa which caused by cocci was the highest, with 86 cases, accounting for 44.10%; 63 cases of otitis externa which caused by fungi and / or Malassezia, accounting for 32.31 %; 31 cases of parasitic otitis externa which caused by mites, accounting for 15.90%; 8 cases of otitis externa which caused by foreign body or IgE (+), accounting for 4.10%. Other causes included auricular hyperplasia and ear tumors in 7 cases, accounting for 3.59%. The causes of different types of dog otitis externa were shown in Figure 2.

**Fig. 2.**
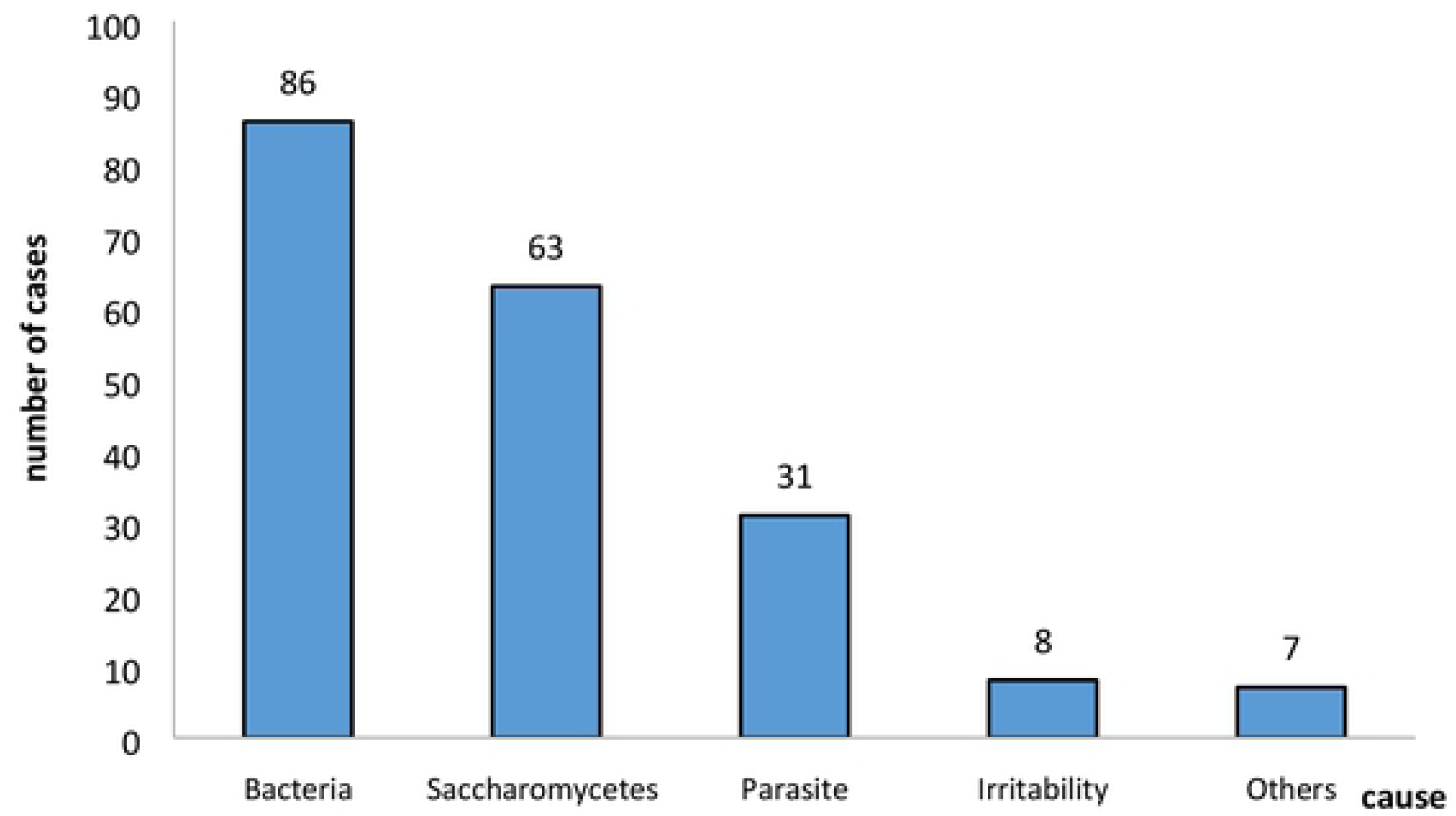
The distribution of causes dog otitis extema from 2012 to 2016

### 2.3 Investigation results of the age of dog ear diseases

This study mainly investigated the incidence of dog ear diseases at three age stages. The statistical results showed that the incidence of ear disease was the highest in elderly dogs (>7 years old), accounting for 39.82%; followed by adult dogs (1-7 years old), accounting for 35.75%; the incidence of young dogs (<1 year old) was 24.43%. The incidence of ear disease at different age stages was shown in Table 2.

**Table 2.**
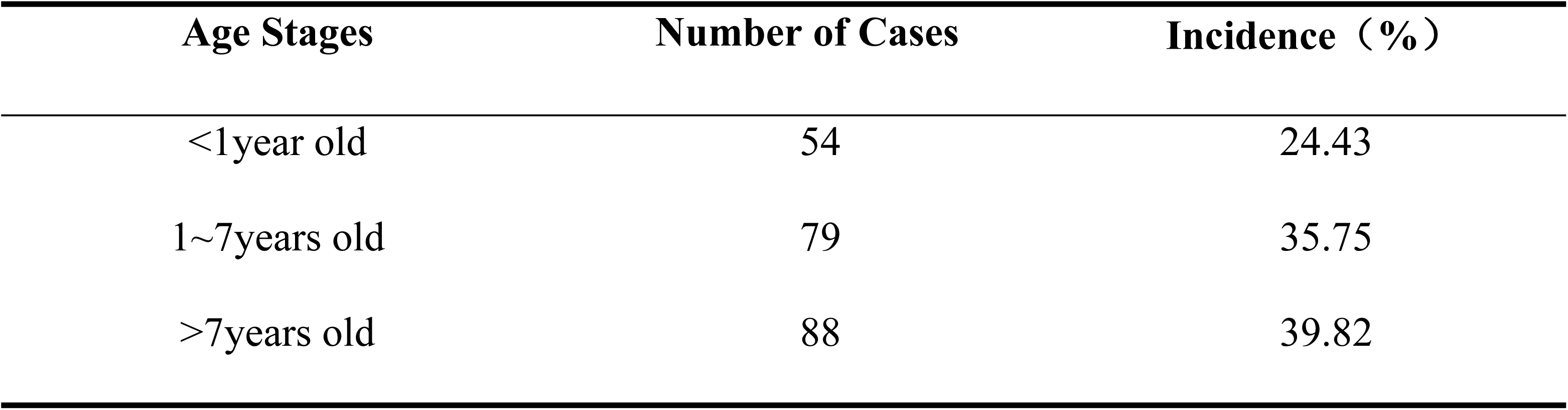
The distribution of dogs at three age stages from 2012 to 2016

Through analysing statistically the incidence of common ear diseases in dogs at three age stages, it was found that the ear diseases of young dogs were mainly otitis externa; the adult and elderly dogs, otitis media, otitis interna, ear skin hyperplasia and ear tumors occur mainly in these two stages. The incidence of common ear diseases in dogs of three age stages were shown in Table 3.

**Table 3.**
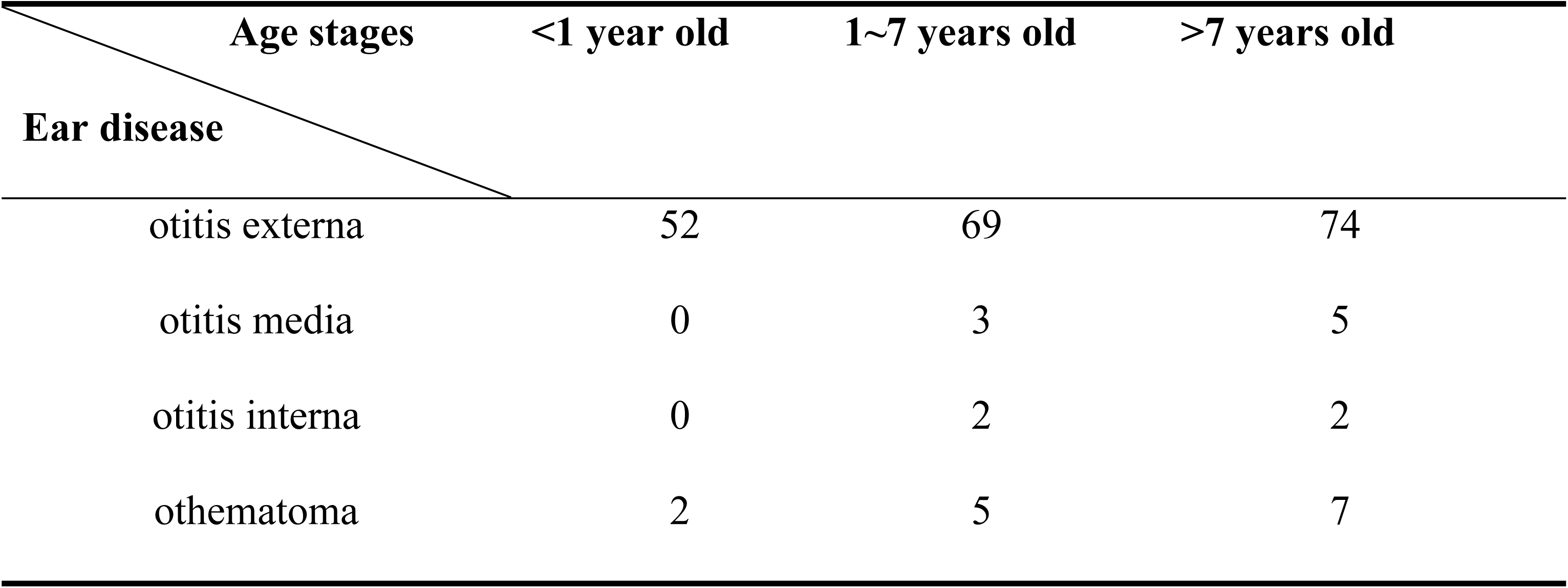
The incidence of common ear diseases in dogs at three age stages from 2012 to 2016

| Age stages<br>Ear disease | <1 year old | 1~7 years old | >7 years old |
| --- | --- | --- | --- |
| otitis externa | 52 | 69 | 74 |
| otitis media | 0 | 3 | 5 |
| otitis interna | 0 | 2 | 2 |
| othematoma | 2 | 5 | 7 |

### 2.4 Investigation results of dog ear diseases in different genders

Among the 221 cases, the number of male cases were 133, accounting for 60.18%; and female cases were 88, accounting for 39.82%. The incidence of canine ears of different genders was shown in Figure 6.

### 2.5 Investigation results of dog ear diseases in different months

In Xi’an, there were multiple cases of dos ear diseases from June to October, with high incidences in August and September, 36 cases and 23 cases respectively, accounting for 16.29% and 10.41% of the proportion of ear diseases. The incidence of dog ear diseases in different months was shown in Figure 4.

### 2.6 Investigation results of dog ear diseases in different breeds

The results of this study had showed that the top three breeds of ear diseases in Xi’an were Teddy, Cocker Spaniel and Golden Retriever, accounting for 18.55%, 10.41%, and 9.50% of the total cases respectively. The incidence of different breeds was shown in Table 4.

**Table 4.** The incidence of dog ear diseases in different breeds from 2012 to 2016

| Breed | Number | Incidence | Breed | Number | Incidence |
| --- | --- | --- | --- | --- | --- |
| Teddy | 41 | 18.55 | Bulldog | 6 | 2.71 |
| Cocker spaniel | 23 | 10.41 | Huskie | 6 | 2.71 |
| Golden retriever | 21 | 9.5 | German Shepherd Dog | 6 | 2.71 |
| Pekingese | 13 | 5.88 | Chow | 3 | 1.36 |
| Schnauzer | 13 | 5.88 | Sharpei | 3 | 1.36 |
| Mix dog | 12 | 5.43 | Rottweiler | 2 | 0.9 |
| Pomeranian | 11 | 4.98 | Chihuahua | 2 | 0.9 |
| Labrador retriever | 10 | 4.52 | Border Collie | 2 | 0.9 |
| Shiba inu | 9 | 4.07 | Yorkshire terrier | 2 | 0.9 |
| Akita | 8 | 3.62 | Bichon Frise | 2 | 0.9 |
| Poodle | 8 | 3.62 | Doberman Pinscher | 2 | 0.9 |
| Samoyed | 7 | 3.17 | Corgi | 2 | 0.9 |
| Shih tzu | 6 | 2.71 | Shelties | 1 | 0.5 |

## 3 Discussion

### 3.1 The epidemiological analysis of the incidence of dog ear diseases

The number of ear diseases showed a tendency to increase with time gradually in Table 1. Firstly, as people’s living standards have improved in recent years, the number of pets has increased gradually, and the number of diseases has also increased. Secondly, with the development of the pet industry in Xi’an, the awareness of the management of some animal owners has strengthened gradually. When animals show symptoms of itching, scratching ears and shaking heads, the owners will bring their pets to the hospital in time.

From 2012 to 2016, the total number of outpatient diseases in Xi’an Teaching Hospital was 20404, with an average number of 4080 cases per year. However, the number of dog ear diseases were less than other diseases, there was 221 in 2012 to 2016, accounting for 1.08%, and the average annual number of dog ear diseases was 44. According to this data, the incidence of ear diseases was still relatively high in the small animal clinic, accounting for more than 10% of the total number of cases, which was inconsistent with the results of this survey. The author thought that there might be the following reasons: Firstly, people had pay great attention to the health of their pets and took effective preventive measures, such as deworming, regular bathing, trimming hair, pulling ears and so on, which would reduce the incidence of the ear diseases to a certain extent. However, some pets’ owners did not have enough awareness of feeding management. They thought that ear diseases were a minor diseases and would not go to the hospital for treatment specifically. Because of other diseases go to the hospital, and the veterinaries will check the ear. Secondly, it may be related to the geographical location and popularity of the hospital. Generally, pets’ owners are used to going to a nearby hospital for the treatment, and when the disease is serious, other veterinaries will recommend them to our hospital. Thirdly, there is a crossover between ear diseases and skin diseases, there are not many cases of ear disease alone, and most of them are co-diagnosed with other cases. Fourth, in recent years, the number of people raising dogs has gradually decreased, and the number of people raising cats and other pets has increased. Dogs are thought to be more active and need walk regularly, which is more destructive. The cat is gentle and quiet, which do not need to walk.

### 3.2 The analysis of the cause of dog ear diseases

From the figure 1 we can find that the highest incidence in dog ear diseases is otitis externa, and the external auditory canal is the structure which is directly in contact with the outside, and it is naturally most susceptible to damage. The external auditory canal is divided into vertical ear canal and horizontal ear canal. This special structure is conducive to the growth and reproduction of pathogenic microorganisms, so the incidence is the highest (Paterson & Tobias, 2012). Followed by ear hematoma, which is mostly developed from otitis externa. When the dog feels ear itchy, scratching excessively can lead to ear hematoma, or the dogs’ temper is fierce, such as German Shepherd, which can easy to fight and bite, these maybe lead to ear hematoma. Otitis media and otitis interna are mainly caused by otitis externa. Generally, only otitis externa is very serious, or repeating episode will affect the structure of the middle ear and inner ear. And when the tympanic membrane ruptures, microbes maybe enter the middle and inner ear, but this case is few.

As can be seen from figure 2, the highest incidence of otitis externa is bacterial infection, and the common bacteria in the ear canal mainly are *Staphylococcus, Pseudomonas* and *Proteus*. In the experiment, the microorganisms in the ear canal of healthy dogs and affected dogs were inoculated into culture medium for microbial culture identification. There were only a few colonies in the plate of healthy dogs, and 383 strains of bacteria were cultured in the affected dog plates, among which the most were *staphylococcus* (199 strains, accounting for 51.96%), the second and third were *corynebacterium* (19.06%) and *streptococcus* (9.14%). Bacteria are ubiquitous in the environment, and there is a normal balance of bacteria in the ear canal. When the environment in the ear canal changes, which is conducive to the reproduction of bacteria (Zhang Di, 2007).

Next is fungal infection, the most common of which is canine *malassezia*. In addition, it has been reported that there are other fungi in the ear canal, such as *Candida, Trichophyton*, etc. (August JR, 1988). Malassezia is a kind of microorganism in the skin and ear canal of humans and animals, which can cause skin disease and otitis externa (Shokri et al., 2010). When there is inflammation in the ear canal, the content of free fatty acids on the surface of the ear canal becomes lower, and the concentration of triglyceride increases, which can provide good condition for the growth of Malassezia. At present, there are 7 species of malassezia such as *Malassezia furfur, Malassczia ovalis,* and *Malassezia globosa,* which have been isolated from the canine ear canal. Among them, the only non-lipophilic fungus is *Malassezia pachydermatis.* so it has strong growth and belongs to the normal flora on canine skin (Dizotti & Coutinho, 2007). In statistical cases, many ear diseases are not single bacterial or fungal infections, but mixed infections. Therefore, in the treatment of drugs must be taken into consideration, not a single treatment.

The third is external parasites, and the main causes of this disease is Otodectes cynotis. Its development cycle includes four stages: egg, larva, nymph and adult. Dogs show clinical symptoms after infecting 2-3 weeks (Zheng Lili, 2007). The mite lives on the surface of the dogs’ ear canal, and does not enter the inner layer of the skin. It uses the mouthparts to pierce the skin to feed, and it secretes toxic substances during feeding, which causes chemical irritation for the epidermis. The dogs show itchy ears. Because the Otodectes cynotis has a hard outer layer and is highly resistant to the environment, its often survives after leaving the host for several months, so the treatment cycle is relatively long and prone to relapse.

The fourth is allergic diseases. The auricle is an extension of the skin on the surface of the body. In the statistical cases, the cause of allergy were mainly foreign body or IgE. When animals are exposed to allergens many times, which stimulates B cells to produce IgE, and antibodies accumulate to a certain extent. When re-contacting the antigen, IgE binds to the antigen, which can cause a series of reactions in the body. Including the skin of the ear canal becomes itchy and swollen. The older the dog is, the more severe the itching is. The otitis externa is the only appearance of an allergic cause. After a rigorous diagnosis, 75% of canine otitis externa is associated with hereditary hypersensitivity, and about 50% show clinical symptoms. 80% of food allergy cases have otitis externa. The only symptom of about 20% of dogs is otitis externa (Xia Zhaofei & Hao Lili, 2003).

### 3.3 The relationship between the incidence of canine ear disease and age

It can be seen from table 2, the collected cases in Xi ‘an Teaching Hospital, the incidence of ear diseases in adult dogs and elderly dogs are higher than that in young dogs. According to the literature, otitis externa can occur in every age stage of dogs, but most are concentrate on 3 to 6 years old (Nascente et al., 2010), which is roughly consistent with the statistical data in this paper. The incidence of otitis externa is very high in each age stage, 26.67% in young dogs, 35.38% in adult dogs and 37.95% in elderly dogs. For young dogs, first of all, the development of organ is not yet fully mature, the barrier function and the immunity of skin is relatively weak. Secondly, young dogs enjoy outdoor activities and they may be exposed to many allergens, such as ear mites, which increases the risk of allergies. For elderly dogs, the secretion of sebum is mainly affected by hormones. The older dogs have lower function and the less hormones it secretes, so the less nutrients that can be provided to the microorganisms, and the environment is not suitable for microbial proliferation (Li Lanjuan, 2002). Therefore, the incidence of elderly dogs should be low. However, due to the organic function of the elderly dogs are gradually degenerating, the immunity of the skin is reduced, which easily lead to the invasion of pathogenic microorganisms, so the incidence is high (Muller et al.,1989). The organic development of adult dogs tend to be mature, it also has high hormone secretion and sebaceous glands are also growing hypofunction in the ear canal, providing a good growth environment for microorganisms. If the epithelial keratinization is poor in the ear canal, the cellular debris can not be cleaned up in time, forming the accumulation of otolith in the ear canal,it can also cause otitis externa.

Otitis media, otitis interna and otitis hematoma mainly occur in adult dogs and elderly dogs. For young dogs otitis media and otitis interna are 0% in the statistical results. The serious condition of otitis externa can lead to otitis media and otitis interna. In young dogs, if the pruritus was found, people will go to the hospital for treatment immediately, The condition is relatively light, so young dogs rarely have otitis media and otitis interna. The methods to keep the external auditory canal clean is mainly through the migratory function of the epithelium (Fraser G, 1965). The sebaceous glands of adult dogs are continuously secreted, the probability of otitis media will increase. Because adult dogs also like to fight, the ear hematomas are more likely to occur. For elderly dogs, the function of cellular keratinization is decreasing, and the incidence of ear canal hyperplasia and ear canal tumors is greatly increasing.

### 3.4 The relationship between the incidence of canine ear disease and gender

Among the collected cases, there were 133 male cases, accounting for 60.18% of the total cases, which was more than 39.82% of females (Fig.3). In the reviewing literature, it is not clear whether there is any direct correlation between the incidence of canine ear disease and gender (Philip DM, 1988). As for the statistical results, the ratio of male to female was 1:1.5, the difference is significant. It may be related to the secretion of hormones. In dogs, the secretion of sebaceous glands is affected by hormones, and the androgen may lead to hyperplasia and hypertrophy of sebaceous glands. The exuberant sebum provides a highly nutritious environment for microorganisms, which is suitable for breeding. However, the estrogen causes degeneration of the sebaceous glands and declines in function, so females may have a lower risk which have ear diseases than males. Therefore, some endocrine diseases may cause abnormal secretion of sebaceous glands, which may cause ear disease. Secondly, the nutritional level of dogs also improves with the development of people’s living standards. The intake of meat is obviously excessive, which leads to the abnormal exuberant secretion of sebaceous glands, thus causing the clinical symptoms such as pruritus and swelling (Huang Di, 2014).

**Fig. 3.**
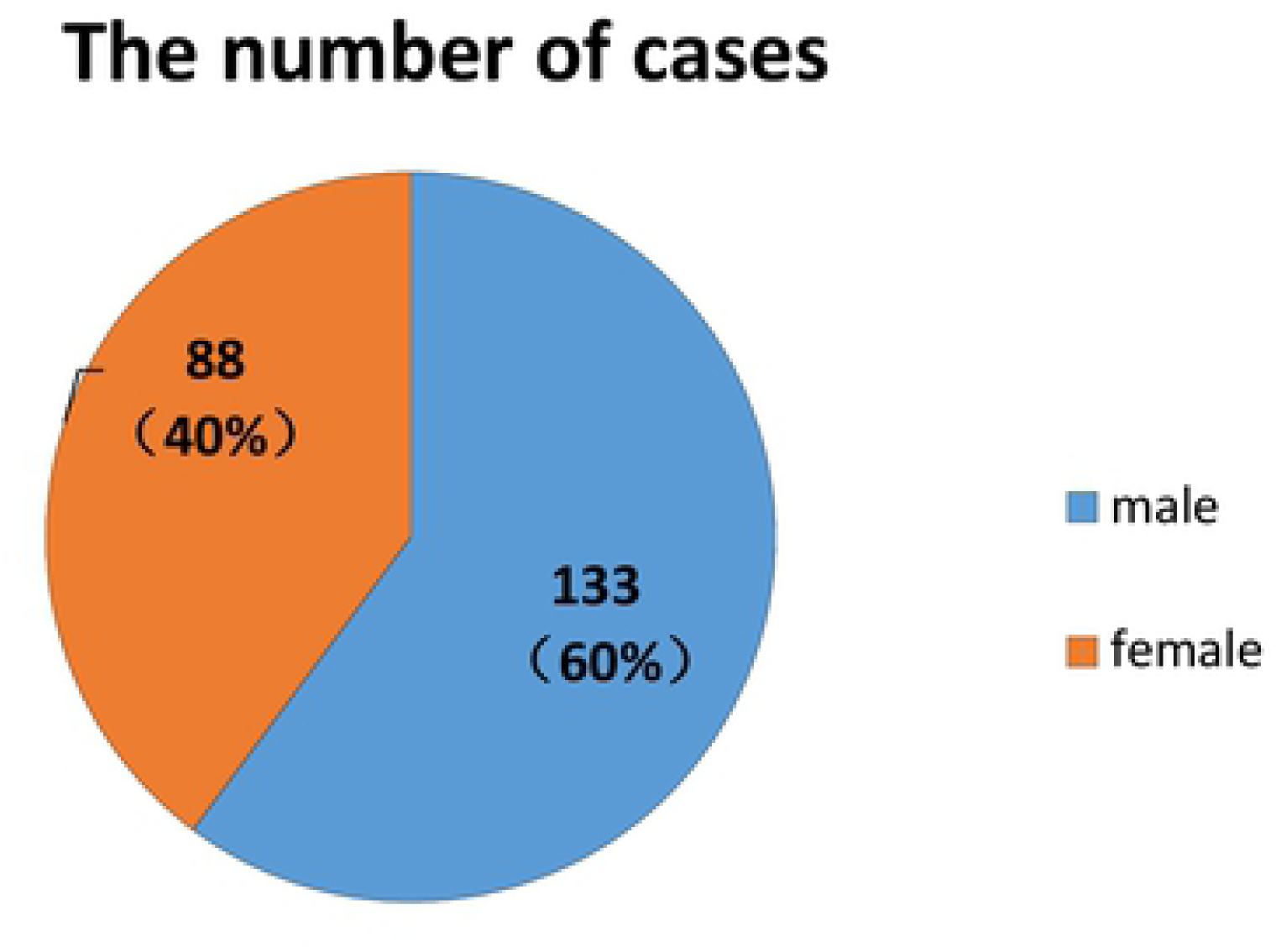
The distribution of dog ear diseases in different genders

**Fig. 4.**
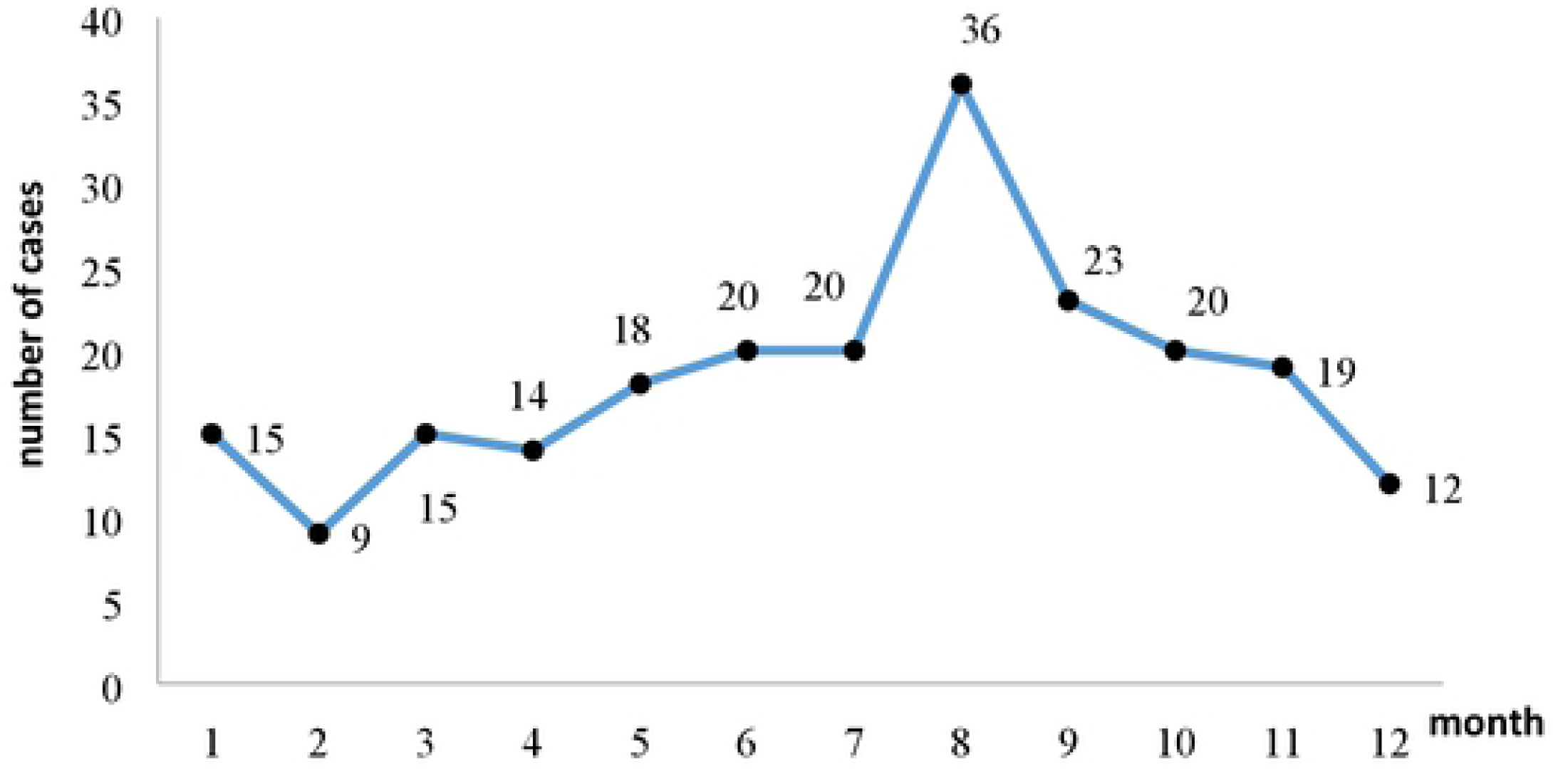
The distribution of dog ear diseases in different months from 2012 to 2016

### 3.5 The relationship between the incidence of canine ear disease and the season

As can be seen from Figure 4, the high incidence of ear diseases in Xi’an is from June to October, accounting for 56.46% of the total. Among them, August and September were high-risk periods, with 36 cases and 23 cases respectively, accounting for 16.29% and 10.41% of the proportion of ear diseases. It can be concluded that the epidemic seasons of ear disease in Xi ‘an are mainly in summer and autumn. Xi ‘an is the biggest city in northwest of China, which has four distinctive seasons, hot and rainy in summer, cool and rainy in autumn. The proportion of precipitation in summer and autumn accounts for more than 70% of annual precipitation. Many literatures have shown that seasonal changes may also lead to the development of ear disease, through the analysis of cases, there is a certain relationship between the incidence and the onset season (Liu Fangyu & Li Yongxue, 2016).

First, Xi’an has a large rainfall in the summer and autumn, with a long rainy period and a high air humidity. However, the structure of the ear canal is relatively closed, and the moist environment is beneficial to the growth of microorganisms, which increases the incidence of bacterial and fungal ear diseases. In addition, Xi’an also has high temperature in summer and autumn, the body surface of dog is covered with hair without sweat glands, so it cannot get rid of the heat through sweat glands, which can only be relieved by mouth breathing. In this way, the capillaries on the body surface of the dog are heated and expanded for a long time, which is conducive to the invasion of parasites (Chen Yi, 2014).

Second, in summer and autumn, dogs spend more time outdoors, and there are more opportunities to be exposed to pathogenic microorganisms. Parasites are more active in summer and autumn, such as ear mite and fleas (Li Zhihong et al., 2016). In addition, Xi’an has a high temperature in summer and autumn, and the owner will increase the number of bathing, which results in dermatic barrier to be broken, the skin is loss of protection, and the risk of illness is increasing significantly.

Third, we generally think that the degree of hair growth in the external auditory canal may have an effect on the temperature of the ear canal, which will increase the temperature of the ear canal, and the higher temperature is conducive to the reproduction of microorganisms. Someone tested 650 dogs with no clinical manifestations of otitis externa and used thermometer to measure the temperature of the external auditory canal. The results showed that the degree of hair growth in the ear canal did not increase the temperature in the ear canal, but decreased. This is inconsistent with our previous assumptions, so there is a dispute on whether the growing level of hair will have an impact on the temperature of the ear canal and lead to ear disease (Huang HP & Huang HM, 1999).

### 3.6 The relationship between the incidence of canine ear diseases and breed

According to the statistical results, cases of ear diseases in teddy dogs are high, accounting for 18.55% of the total. First of all, the teddy dog are cute and popular among young people in northwest of China, so the number of breeding has increased year by year. Then the number of cases will increase accordingly. Secondly, the breed has more hair on the ear canal, which prevents air circulation and keeps the ear canal in a humid environment for a long time. It is conducive to microbial reproduction and easy to cause otitis externa. Some scholars have pointed out in the research report that dogs with drooping auricles are more likely to get ear diseases if they do not consider the density of hair in the ear canal as a pathogenic factor (Logas DB, 1994). In second and third place are cocker spaniels and golden retrievers, which accounted for 10.41% and 9.50% of the cases, respectively. Both the cocker and the golden retriever are medium and large dogs. The hair in the ear canal is more than other breeds, and they also have large and drooping auricles. Therefore, comparing with the dogs with vertical auricle, the drooping auricle prevents the air circulation and heat dissipation in the ear, which results in a moist environment in the ear canal. It is conducive to make the reproduction of microorganisms. And the cerumen glands in their ear canal are rich. Under the stimulating of bacteria or parasites, the cerumen glands produce a large amount of substances, and the exuberant hair causes the secretions to remain mostly in the ear canal, which can provide abundant nutrients for microorganisms to grow and reproduce.

However, there are also some breeds with drooping auricles such as beagle dogs, which have little incidence in developing ear diseases. Some species with vertical auricles such as huskies have higher incidence. Therefore, after analyzing, it is consider whether auricle droops or not, it is not the direct cause of otitis externa, but it may affect the therapeutic effect. In addition, ear anatomy of some breeds is different, that also can cause ear diseases. In some breeds, such as sharkskin and bulldog, the structure of ear canal is congenitally long and narrow and it has a certain angle to the horizontal plane, which makes the secretion be not discharged normally, increasing the incidence of ear diseases. Some are acquired, such as ear canal edema, ear canal mass and ear canal hyperplasia, which may press the ear canal, resulting in stenosis of the ear canal and causing ear diseases.

In different regions, the results of the survey are different. This may be related to the following aspects. Firstly, it is related to the habits of local residents. For example, there is a large number of Beijing dogs in Beijing before, so the incidence of Beijing dogs is higher in Beijing. In recent years, the number of Teddy, Pomeranian, and Golden Retriever in Xi’an has increased, so the incidence has also increased greatly. Secondly, some hairy breeds, such as schnauzer and teddy, which get regular haircut. When plucking hair of ear canal, the external force will destroy the stratum corneum of the ear canal skin, and the defense of skin is weakened, it is prone to cause ear diseases.

## Acknowledgments

We are grateful to all of those who helped with this study, especially to hospital staff of the Xi’an Teaching Hospital of Northwest A&F University. This study was supported by the Fundamental Research Funds for the Central Universities (Grant No. 2452016039), the Natural Science Foundation of Shaanxi Province of China (Grant No. 2014JQ3086) Foundation for Talent of Northwest A&F University (Grant No. Z109021110).

## Reference

Chen Yi. Epidemiologic investigation and control strategy of canine dermatonosis in taizhou city[D]. Yangling: Northwest A&F University, 2014.

Dizotti CE, Coutinho DA. Isolation of Malassezia Pachydermatis and M.sympodialis from the External Ear Canal of Cats with and without otitis External[J]. Acta Veterinaria Hungarica, 2007, 55(4): 471–477.

Fraser G. Aetiology of Otitis Externa in the Dog[J]. Small anim Pract, 1965, 6: 445–452.

HP Huang, HM Huang. Effects of ear type, sex, age, body weight, and climate on temperatures in the external acoustic meatus of dogs[J]. Am J Vet Res, 1999, 60: 1173-1176.

Huang Di. Evaluation of drug resistance to aminoglycosides in pathogenic microorganism of canine otitis externa[D]. Beijing: China Agricultural University, 2014.

August JR. Otitis externa-a disease of multifatorial etiology[J]. Vet Clin Small Anim, 1988, 7(4): 731–742.

Li Lanjuan. Infectious Microecology, 2rd Editoin[M]. Beijing: People’s Medical Publishing House, 2002: 486–491.

Li Zhihong, Li Zhiqi, Qi Yingying, Wangjing. Diagnosis and treatment of Otodectes canis[J]. Contemporary Animal Husbandry, 2015, 2: 18–19.

Liu Fangyu, Li Yongxue. The relationship between the incidence of canine skin disease and the season[J]. Heilongjiang animal science and veterinary medicine, 2016, 03: 147–148.

Logas DB. Diseases of the ear canal[J]. Vet Clin North Am Small Anim Pract, 1994, 24(5): 905–919.

Muller GH, Kirk RW, Scott DW. Small Animal Dermatology[M]. Philadelphia: WBS aunders, 1989: 718–721, 807-809, 811.

Nascente PDS, Santin R, Meinerz ARM, et al. Estudo da Frequencia de Malassezia Pachydermatis em Caes Com Otite Externa no Rio Grande do Sul[J]. Ci Anim Bras, 2010, 11(4): 527–536.

Paterson S, Tobias K. Atlas of Ear Diseases of the Dog and Cat[M]. Wiley-Blackwell, 2012: 1–45.

Philip DM. Preventive ear care for dogs and cats[J]. Vet Clin North Am Small Anim Pract, 1988, 18(4): 845–858.

Shokri H, Khosravi A, Rad M, et al. Occurrence of malassezia species in Persian and domestic short hair cats with and without otitis externa[J]. Journal of Veterinary Medical Science, 2010, 72(3): 293–296.

Xia Zhaofei, Hao Lili. The cause of otitis in dogs and cats[J]. Chinese Journal of Veterinary Medicine, 2003, 39: 44–45.

Zhang Di. A retrospective study on clinical epidemic characteristics and identification of microorganism in the external ear canal and evauation of susceptibility testing in vitro of Otitis Externa in canine[D]. Beijing: China Agricultural University, 2007.

Zheng Lili. Prevention and treatment of urticaria mite in dogs and cats[J]. Chinese Journal of Animal Husbandry and Veterinary Medicine, 2007, 9: 92.

